# Results dissemination from clinical trials conducted at German university medical centres was delayed and incomplete

**DOI:** 10.1101/467746

**Authors:** Susanne Wieschowski, Nico Riedel, Katharina Wollmann, Hannes Kahrass, Stephanie Müller-Ohlraun, Christopher Schürmann, Sean Kelley, Ute Kszuk, Bob Siegerink, Ulrich Dirnagl, Jörg Meerpohl, Daniel Strech

## Abstract

**Objective:** Timely and comprehensive reporting of clinical trial results build the backbone of evidence-based medicine and responsible research. The proportion of timely disseminated trial results can inform alternative national and international benchmarking of university medical centers (UMCs).

**Study Design and Setting:** For all German UMCs we tracked all registered trials completed between 2009 and 2013. The results and an interactive website benchmark German UMCs regarding their performance in results dissemination.

**Results:** We identified and tracked 2,132 clinical trials. For 1,509 trials, one of the German UMCs took the academic lead. Of these 1,509 “lead trials”, 39% published their results (mostly via journal publications) in a timely manner (<24 months after completion). More than six years after study completion, 26% of all eligible lead trials still had not disseminated results.

**Conclusion:** Despite substantial attention from many stakeholders to the topic, there is still a strong delay or even absence of results dissemination for many trials. German UMCs have several opportunities to improve this situation. Further research should evaluate whether and how a transparent benchmarking of UMC performance in results dissemination helps to increase value and reduce waste in medical research.

## Background

The results of clinical trials build the backbone of evidence-based medicine. They inform clinical decision making [1] and health technology assessment [2, 3]. They also inform decision making within ongoing trials and decision making related to the design, review, and funding of new trials [4]. Non-dissemination or delayed dissemination of trial results negatively affects all of these decisionmaking processes [5–8]. For three decades studies investigated and criticized this challenge [9–11]. Since 2008, the Declaration of Helsinki includes the requirements that every study involving human subjects should be prospectively registered and that all results should be made publicly available – irrespective of the results’ direction [12]. The joint statement by the World Health Organization in 2015 defined “timely publication” as “24 months for publication in a peer-reviewed journal (preferably open-access) and 12 months for publication of the key results in the registry’s result section” [8].

The current dissemination of clinical trial results still looks quite different. Recently, Chen et al. analysed the publication of more than 4,000 interventional clinical trials across all 51 US university medical centres (UMCs) that were completed between October 2007 and September 2010 [13]. Only 29% of trials published their results within 24 months after study completion, and only 13% of trials posted their summary results (a tabular summary of the key outcomes) on the clinical trials registry ClinicalTrials.gov. Overall, as of July 2014, 35% of all trials were found to be unpublished [13]. Schmucker et al. conducted a systematic review on similar tracking investigations with a total of 5,112 studies and found that on average 54% of studies are unreported, with a range from 24% - 74% for the individual tracking investigations [14]. A similar range was detected in the systematic review of Dwan et al. [15].

In recent years, journals [16], agencies (the Food and Drug Administration (FDA) and European Medicines Agency (EMA)) [17], ethical guidelines [18], and most recently, funding bodies [19] have all explicitly highlighted the need to reduce biased or delayed publication and developed policies to proactively achieve this objective. UMCs, in contrast, which function as trial sites and host the responsible principle investigators (PIs), have remained surprisingly silent about this issue [20, 21]. A transparent benchmarking for how complete and timely UMCs are in reporting their trial results could incentivize the implementation of more effective UMC policies in this regard. Such benchmarks could also raise public and media awareness about this issue. The “TrialsTracker” project, which includes several different trackers provides automatically updated data for benchmarking activities [22, 23]. While the FDA (https://fdaaa.trialstracker.net) and EU (http://eu.trialstracker.net) TrialsTrackers check if summary results were posted on the ClinicalTrials.gov and the EU clinical trials register (EUCTR), respectively, the original TrialsTracker (https://trialstracker.ebmdatalab.net/) additionally searched for linked results on Pubmed (last update March 2017). TrialsTracker increases its public outreach by presenting results via a publicly accessible website. The FDA and EU TrialsTrackers, however, have two limitations. First, they focus on trials that fall under mandatory reporting rules according to the FDA Amendment Act (FDAAA) or the European Commission guideline 2012/c302/03. Second, their method is restricted to the automated search of registry entries.

In this study, we further develop the concept and practice of benchmarking UMCs in three ways. First, with regard to the sample, we sampled and followed up trials (i.e. all registered interventional clinical studies) that had a completion date between 2009 and 2013 for all German UMCs. In a continuously updated map of all studies on ClinicalTrials.gov Germany is second with 17,945 trials behind France with 21,423 trials [24]. According to the advanced search of the EUCTR Germany has most trials with a EudraCT protocol (n=10,273) followed by United Kingdom (n=8,589), Spain (n=8,536), Italy (n=7,057) and France (n=4,551) [25]. Second, we extended the standard publication search strategies to comprehensive hand searches in Google Scholar. This allow a better understanding of the full picture of available results published outside registry websites and PubMed. Third, with regard to benchmarking, we developed a website, including a Shiny app, that allows the interactive visualization of benchmarking according to the different variables that influence publication measurement, such as time to publication, publication format, sponsor, timing of registration, completion date and others.

## Methods

The protocol for this project, including all methodological details for sampling and following up clinical trials for data extraction, and statistical analyses was preregistered with the Open Science Framework (OSF) and continuously updated for amendments (https://osf.io/fh426/). In the following sections, we summarize the methods.

### Retrieval of trials

We downloaded the AACT dataset, which aggregates information from ClinicalTrials.gov into a relational database, from http://aact.ctti-clinicaltrials.org/ (version date: April 17, 2017). We further downloaded the dataset from the German Clinical Trials Registry (DRKS) from www.drks.de on July 27, 2017. The delay in DRKS extraction was due to a slight change in the filtering criteria (see detailed methods on OSF). We used an R script to combine all relevant datasets and to extract the trial characteristics.

### Inclusion and exclusion of studies

R was also used to restrict the resulting dataset to studies with a primary completion date (AACT, can be planned or actual) or study end date (DRKS, actual) in the years 2009-2013, as well as to exclude observational studies, incomplete entries (missing NCT, affiliation or primary completion date), and duplicates.

For all studies we checked if one or several of the German UMCs contributed to the trial by searching for different versions of the UMC names in the affiliation fields of the extracted dataset. Contributions of a UMC to a trial were either counted as a) “lead” contribution, where the UMC had a mention as responsible party, lead sponsor or principal investigator (AACT) or as primary sponsor (DRKS), or b) “facility” contribution, where the UMC only recruited patients as a facility or collaborator (AACT) or recruitment location (DRKS). One trial can be counted for multiple UMCs if they have different contributions to the trial. However, for the overall results reported in this study, each trial associated to several UMCs is counted only once. After automatic filtering for the UMC names, the correct assignment of trials to UMCs was verified manually.

In the following, we concentrate on the results for the lead trials only. For further results regarding the facility trials see http://s-quest.bihealth.org/intovalue/.

Only studies with the status “Completed”, “Terminated”, “Suspended”, or “Unknown” (or the equivalent DRKS categories; see detailed methods on OSF) were included. Studies from the DRKS sample that also appeared in the AACT sample were identified by searching for NCT IDs as secondary IDs in the DRKS dataset and subsequently removed.

### Publication search

For each of the included studies, a results publication was searched independently by two researchers in a 3-step process between July 2017 and December 2017 (see also Figure 1, search strategy). 1) The clinical trial identifier (NCT ID or DRKS ID) was entered on ClinicalTrials.gov/DRKS.de, and the earliest result publication linked in the registry was searched and checked if it was indeed a results publication for the trial. Reviews and other background literature were excluded. 2) The clinical trial identifier was entered on PubMed. 3) Google Scholar and (if no hit was found) Web of Science were searched by subsequently entering the following search terms: clinical trial identifier, official title, brief title (if available), intervention name and principal investigator or primary sponsor. The first two results pages were screened. Publications were matched using a list of explicit criteria (i.e., study design, intervention, and outcomes). All criteria needed to be met to be counted as a match.

**Figure 1:**
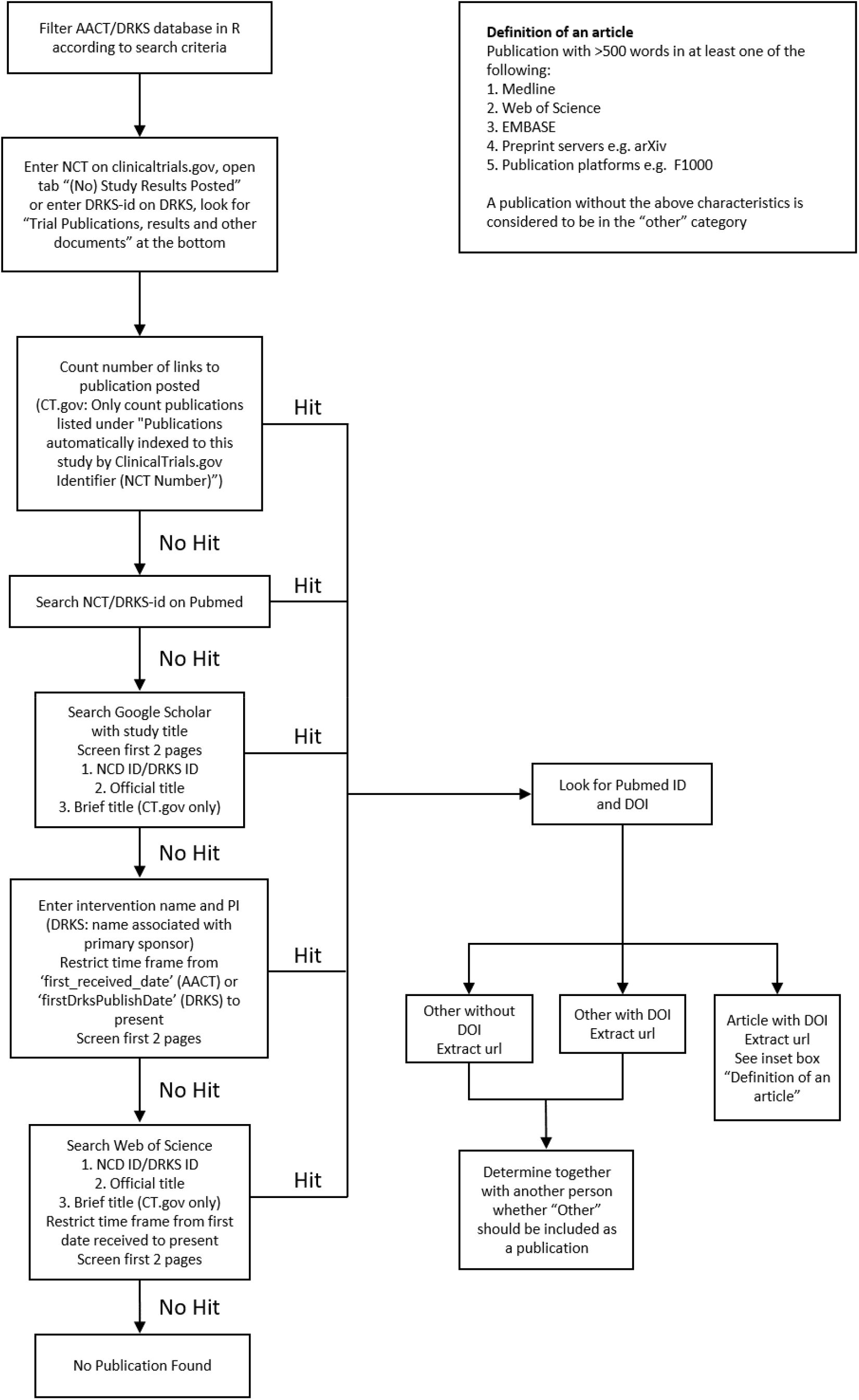
Search strategy

If, after all three searches, there was still no result, the study was characterized as “no publication found”. Additionally, the researchers checked if a summary result was posted on clinicaltrials.gov.

In some cases, we identified results resources that were not journal publications (“other” results category). We counted doctoral theses containing the trial results as results publications, but we did not count conference abstracts, posters or presentation slides.

### Interrater reliability

For all studies two researchers independently conducted the publication search. Both researchers compared the independently identified results publications. If they identified different publications, we counted the earliest publication. Interrater reliability, which was defined as how often two raters had independently identified the same results publication, was at 78%. In cases with different publications we reached a 100% agreement on which is the earlier publication.

For further information on interrater reliability, data extraction, R-scripts, and statistics (logistic regression and Kaplan-Meier), see the abovementioned protocol registered with OSF (https://osf.io/fh426/).

## Results

### Demographic data

We identified 2,132 clinical trials via clinicaltrial.gov (n=1,905) and DRKS (n=227) that i) recruited trial participants from at least one German UMC and ii) had their primary completion date (PCD, last visit of last patient for a primary outcome measure) between 2009 and 2013. These trials included 506,876 anticipated participants.

Altogether, 71% (n=1,457) of all trials were counted as lead trial for one of the corresponding German UMCs. Of these 1,457 lead trials, 502 (35%) investigated drugs, and 266 (18%) investigated devices; the rest were “behavioural”, “procedure” or “other” interventions. Only a minority of these lead trials (n=360; 25%) were registered prospectively, and 878 (60%) were registered more than 21 days after the given start date (can be actual or planned, see supplemental Table 1 on OSF) of the trial, with 246 (17%) registered after the completion date (CD). 113 lead trials (8%) included more than 500 anticipated participants. A total of 1,054 trials (72%) were completed, and 136 (9%) were either terminated early or suspended; for 267 trials (18%), the status was unknown. For additional demographic data, see Table 1.

**Table 1:**
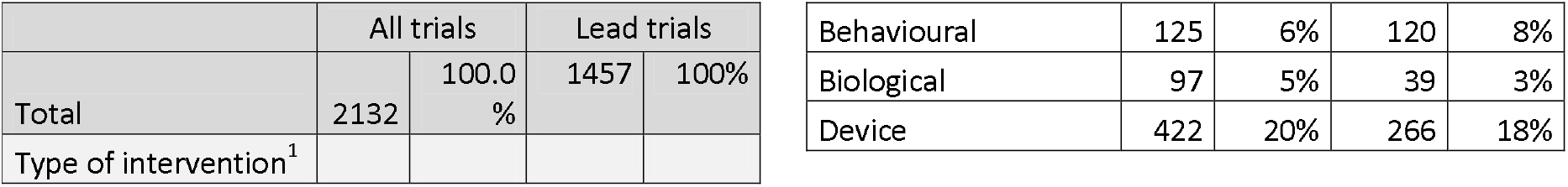
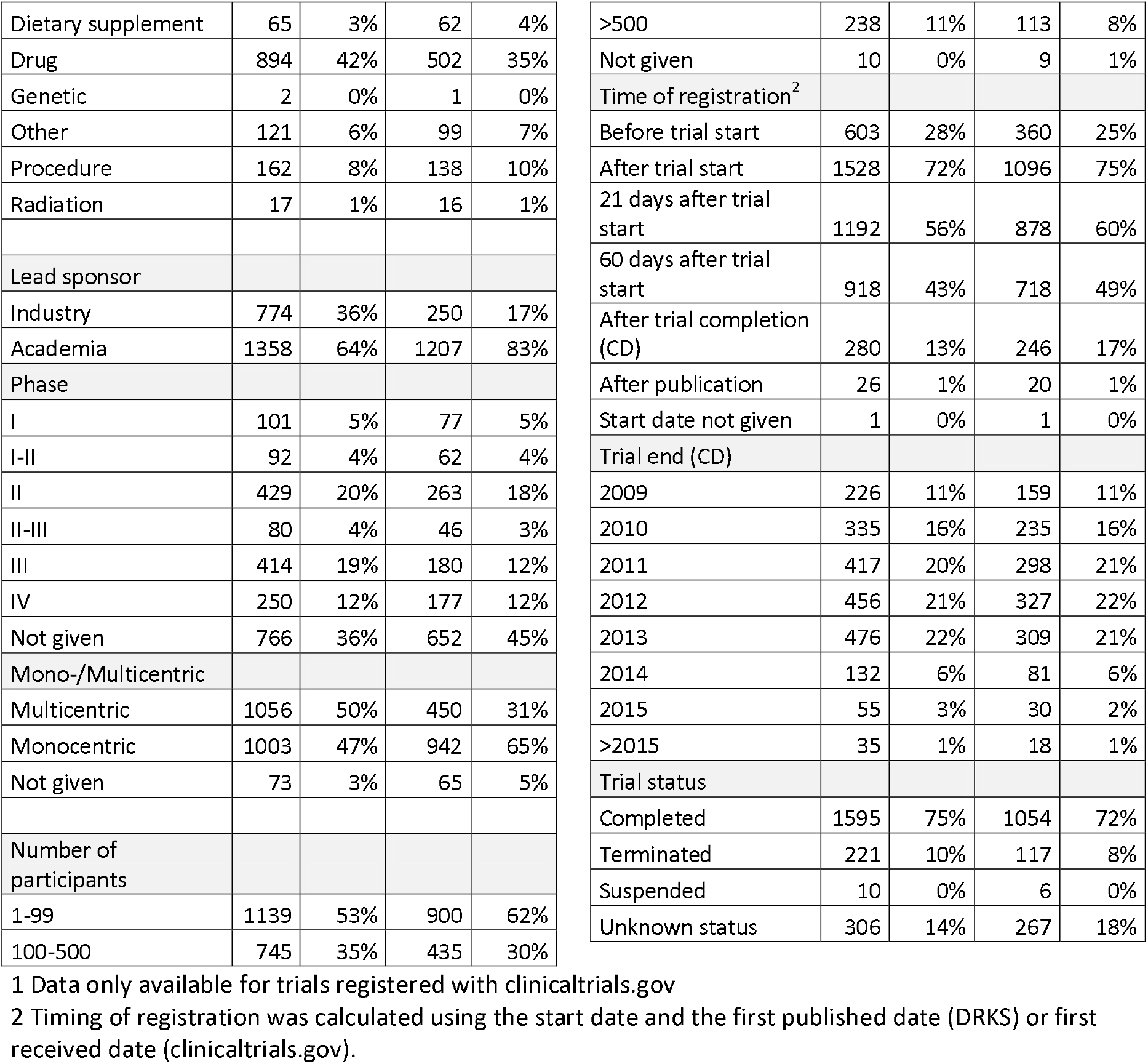
Demographic data for “all trials”

### Overall results reporting and the Shiny app website

Because our paper cannot report on all measurement variables that were applied in our study, we developed an interactive website (based on a Shiny app) that allows users to select and combine the measurement variables in which they are most interested and develop a corresponding benchmark for all 36 German UMCs. The website is http://s-quest.bihealth.org/intovalue/. In the following sections, we report the most essential findings of our study.

Of all 1,457 lead trials, we could follow up 1,438 for a minimum of 24 months after the CD. Of those trials, 39% published their results via journal publications, summary results or dissertations within 24 months after the CD. At the level of German UMCs, this publication rate varied from 14% to 62% (Table 2). We found a steady improvement in timely publication from 35% for trials completed in 2009/2010 up to 42% for trials completed in 2012/2013. Figure 2 presents the percentage of unpublished trials over time. By April 2017, summary results were reported in the registry for 91 (7%) of 1,243 lead trials registered with clinicaltrials.gov. We further identified six dissertations, 43 abstracts (among them conference abstracts), and one presentation.

**Table 2:**
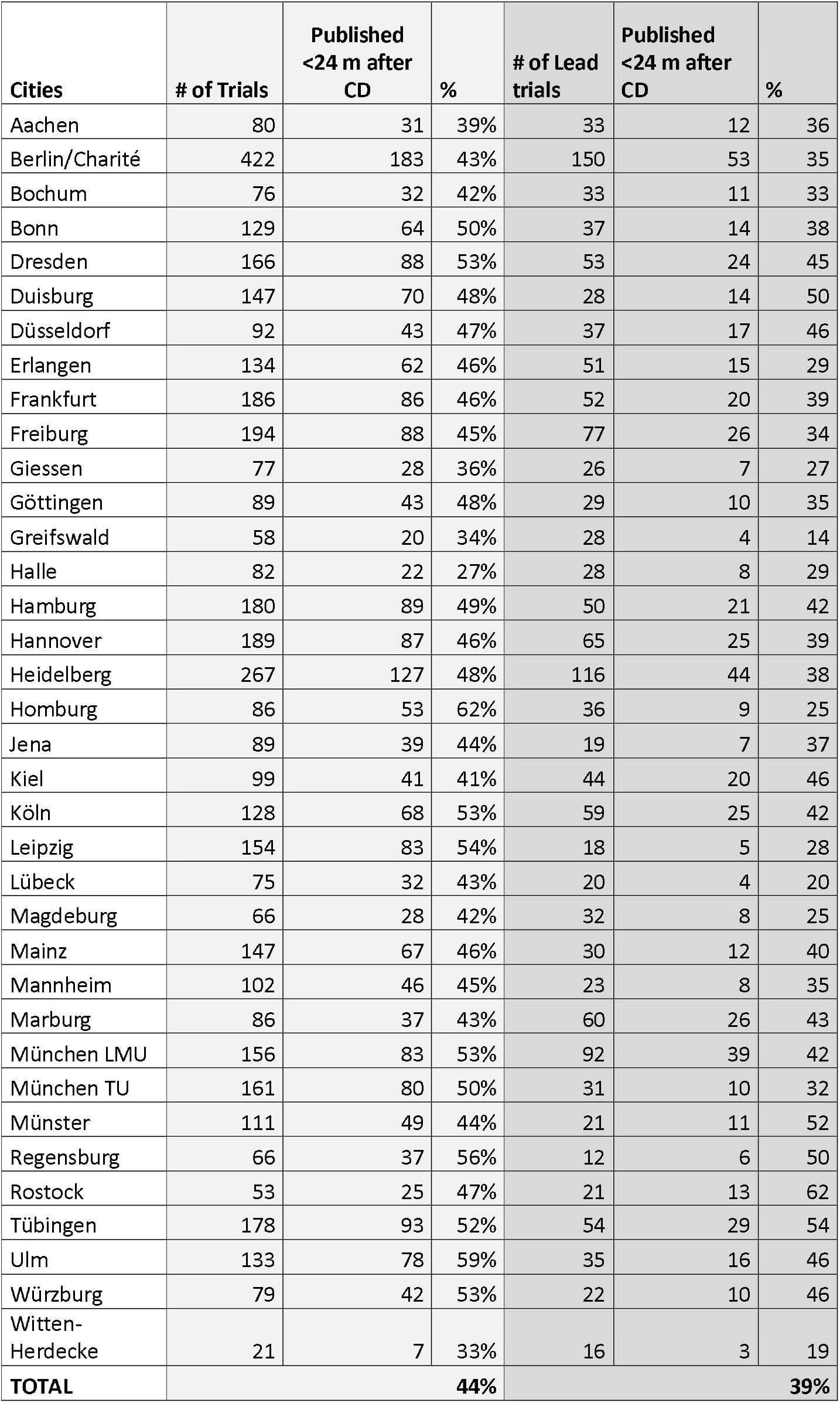
Publication rates at the level of individual German university medical centres (UMCs) → More variations of this table are available on the interactive website http://bit.ly/intovalue.

| Cities | # of Trials | Published<br><24 m after<br>CD | % | # of Lead<br>trials | Published<br><24 m after<br>CD | % |
| --- | --- | --- | --- | --- | --- | --- |
| Aachen | 80 | 31 | 39% | 33 | 12 | 36 |
| Berlin/Charité | 422 | 183 | 43% | 150 | 53 | 35 |
| Bochum | 76 | 32 | 42% | 33 | 11 | 33 |
| Bonn | 129 | 64 | 50% | 37 | 14 | 38 |
| Dresden | 166 | 88 | 53% | 53 | 24 | 45 |
| Duisburg | 147 | 70 | 48% | 28 | 14 | 50 |
| Düsseldorf | 92 | 43 | 47% | 37 | 17 | 46 |
| Erlangen | 134 | 62 | 46% | 51 | 15 | 29 |
| Frankfurt | 186 | 86 | 46% | 52 | 20 | 39 |
| Freiburg | 194 | 88 | 45% | 77 | 26 | 34 |
| Giessen | 77 | 28 | 36% | 26 | 7 | 27 |
| Göttingen | 89 | 43 | 48% | 29 | 10 | 35 |
| Greifswald | 58 | 20 | 34% | 28 | 4 | 14 |
| Halle | 82 | 22 | 27% | 28 | 8 | 29 |
| Hamburg | 180 | 89 | 49% | 50 | 21 | 42 |
| Hannover | 189 | 87 | 46% | 65 | 25 | 39 |
| Heidelberg | 267 | 127 | 48% | 116 | 44 | 38 |
| Homburg | 86 | 53 | 62% | 36 | 9 | 25 |
| Jena | 89 | 39 | 44% | 19 | 7 | 37 |
| Kiel | 99 | 41 | 41% | 44 | 20 | 46 |
| Köln | 128 | 68 | 53% | 59 | 25 | 42 |
| Leipzig | 154 | 83 | 54% | 18 | 5 | 28 |
| Lübeck | 75 | 32 | 43% | 20 | 4 | 20 |
| Magdeburg | 66 | 28 | 42% | 32 | 8 | 25 |
| Mainz | 147 | 67 | 46% | 30 | 12 | 40 |
| Mannheim | 102 | 46 | 45% | 23 | 8 | 35 |
| Marburg | 86 | 37 | 43% | 60 | 26 | 43 |
| München LMU | 156 | 83 | 53% | 92 | 39 | 42 |
| München TU | 161 | 80 | 50% | 31 | 10 | 32 |
| Münster | 111 | 49 | 44% | 21 | 11 | 52 |
| Regensburg | 66 | 37 | 56% | 12 | 6 | 50 |
| Rostock | 53 | 25 | 47% | 21 | 13 | 62 |
| Tübingen | 178 | 93 | 52% | 54 | 29 | 54 |
| Ulm | 133 | 78 | 59% | 35 | 16 | 46 |
| Würzburg | 79 | 42 | 53% | 22 | 10 | 46 |
| Witten-Herdecke | 21 | 7 | 33% | 16 | 3 | 19 |
| **TOTAL** |  |  | **44%** |  |  | **39%** |

**Figure 2:**
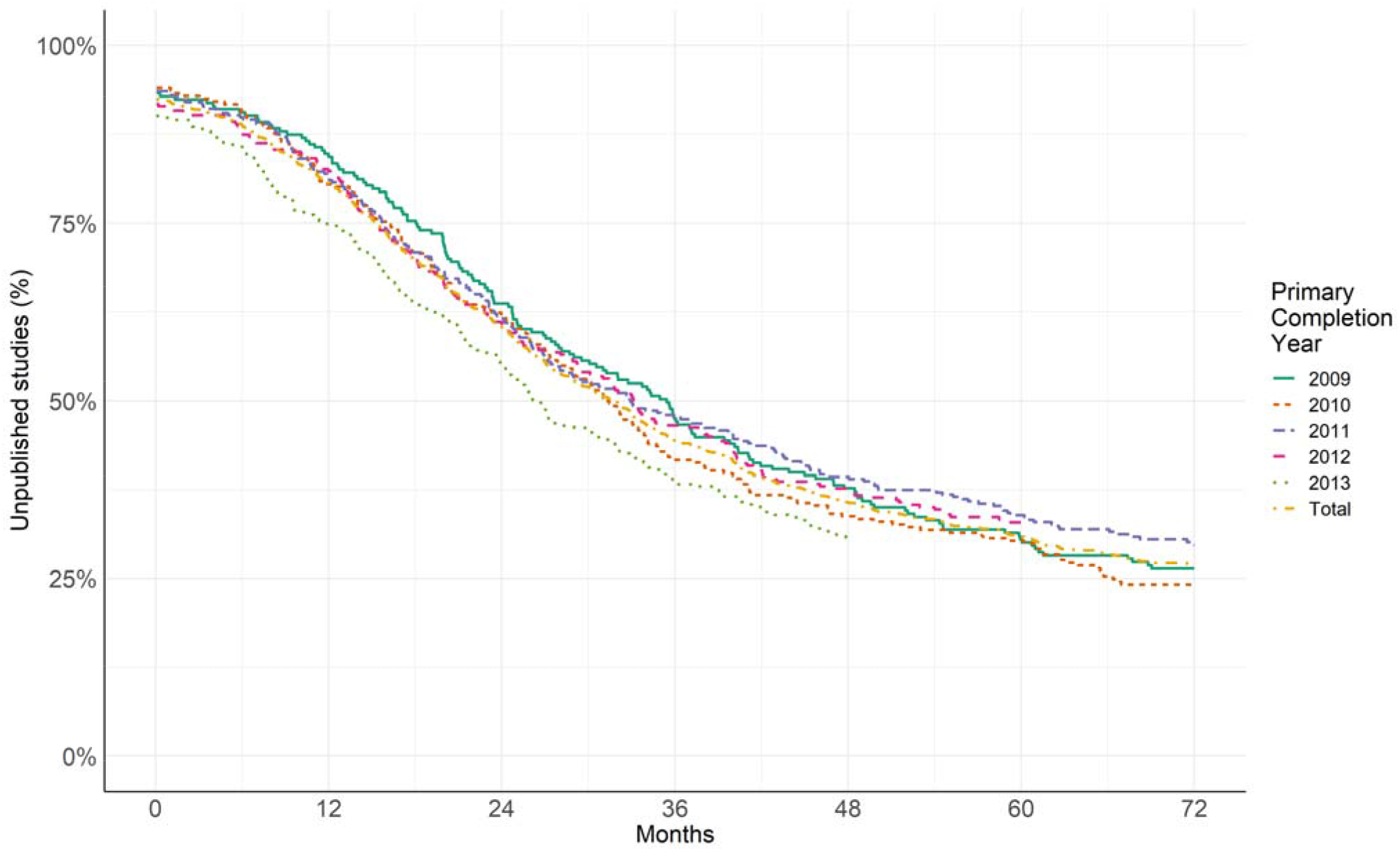
Kaplan-Meier curve showing the percentage of unpublished lead studies over time grouped by primary completion year. Full methods and details under https://osf.io/fh426/. More variations of this graph are available on the interactive website http://s-quest.bihealth.org/intovalue/

Of the 1,457 lead trials, there was a subgroup of 651 trials that we could follow up for more than six years after the CD. For this subgroup, we found an overall publication rate of 74%, with a variation across universities of 56% to 100%. Altogether, 18,305 participants were planned to be included in the 171 trials that have not published their results. Extrapolated to the full sample of lead trials (18,305 × 1,457 ÷ 651 ÷ 5 years) an average of 8,194 planned participants per year were included in trials from German UMCs that did not disseminate their results after more than six years.

All the results presented above were generated by time-intensive searches, including searches in Google Scholar and Web of Science (see Methods), that were performed independently by two researchers with training in literature searching. When restricting our search efforts to more convenient standards (registry and PubMed; see Methods), we could identify results for only 26% of trials within 24 months after CD (vs. 39% with our extensive search) and for 45% of trials followed up for more than six years (vs. 74% with our extensive search). Thus, we could identify 33% of all timely publications and 39% of all publications with a six-year follow-up period only via the additional search strategies.

### Subgroup analyses

The overall publication rates (for more than six years after CD) differed substantially (more than 10%) according to the following factors:

- number of participants (71% for trials with 1-100 participants but 94% for trials with >500 participants),
- timing of registration (69% for prospectively registered trials but 86% for trials registered after the CD), and
- trial status (43% for trials with terminated/suspended/unknown status but 83% for completed trials).

To identify subgroups with substantially different publication rates, we performed an additional exploratory logistic regression analysis (as preregistered, see supplementary file on OSF for more details). We identified the variables ‘mono-vs multicentric’ (OR 1.66, CI 1.39-1.98), ‘lead sponsor’ (industry vs academia, OR 1.67, CI 1.40-2.01), and ‘number of participants’ (OR 1.19 per 500 participants, CI 1.08-1.32) as the variables with the strongest association with publication rates. However, these associations in itself, or in combination are too weak to predict which future studies will be reported in time with great confidence.

## Discussion

In this study, we demonstrate that only 39% of all registered clinical trials conducted at one of the 36 German UMCs published their results in a timely manner within 24 months after the trial’s completion date (CD). This rate further decreases to 26% when applying standard search strategies. Six years after the CD and with the most extensive search strategies, 26% of all trials still remain unpublished.

For the following reasons, this high proportion of delayed or omitted result dissemination is unethical and a substantial waste of important research resources. First, the fact that 26% of all clinical trials withhold the knowledge they gained or delay its dissemination negatively impacts i) the design of future, non-redundant translational research and ii) patient-oriented, evidence-based medical decision making. Second, every year, more than 8,000 participants on average were included in lead trials from German UMCs that did not generate any knowledge gain and thus no social value. Social value, however, is the basic ethical principle justifying research that adds burdens and risks to participants. Moreover, most trial participants are patients who already suffer from a disease. Third, administrative efforts to report summary results in the tabular format required by ClinicalTrials.gov are minimal, and this type of results reporting does not prevent more detailed and contextualized result publications in peer-reviewed journals [26]. Despite the ethical rationale only 7% (n=91) of clinical trials conducted at German UMCs (“lead trials”) reported their summary results in clinicaltrials.gov. The recently published EU TrialsTracker that evaluated the compliance with summary results reporting in the EUCTR) confirmed these low reporting rates [22].

In contrast to most other trial tracking activities our search strategy included additional hand searches in Google Scholar that identified many publications that were not indexed at clinicaltrials.gov or PubMed. This step is critical to increase the number of identified publications. Unfortunately, this time-intensive approach only allows static analysis. If, in the future, results information provided at the registry are more complete (as either summary results or linked journal publications), automated tracking could fully take over.

German UMCs have many unique possibilities to improve the current situation. First, UMC staff that coordinates clinical trials could remind and support PIs in timely reporting. UK universities are currently leading the way in this regard [27]. A second option would be to reward those PIs who manage to publish their results in a timely manner and/or report summary results in the registry. At German UMCs, the performance-oriented allocation of funds (“LOM/Leistungsorientierte Mittelvergabe”) currently only rewards aggregated impact factors and third-party funding. A third and harsher option would be to sanction those PIs who do not manage to report at least summary results in the registry within 24 months after CD. A recent update of the Wellcome Trust funding policy for clinical trials demonstrated that funders at least might decide to go this way [28].

While we applied extensive automated methods and hand searches to track each registered trial by two independent researchers our study has still several limitations. Despite the fact that we identified higher publication rates than all former tracking studies our results might still underestimate the true publication rates because we did not search in scientific databases other than PubMed and Web of Science, and we did not contact the responsible parties. Our results might also overestimate the true publications rates for several reasons. First, most included trials (60%) were retrospectively registered. These trials had substantially higher publication rates, which might reflect a registration and reporting bias. Furthermore, we did not include observational clinical studies in our sample. Former tracking studies that sampled at the level of German institutional review boards (IRBs) reported substantially lower publication rates for observational studies [29]. Additionally, our results might overestimate the time to publication because we stopped searching for further publications once we found the first results publication, possibly missing earlier results publications. We also rely on the registry entries being correct. Entries in the DRKS database might be duplicates of entries in clinicaltrials.gov without an appropriate cross-referencing of the NCT ID. Completion dates are entered as expected dates and might not always reflect the actual completion dates. As registries are increasingly consulted as a key resource for health care information and for quality assessment of research practices it is important to improve their quality, accuracy and timeliness [30]. We published a more detailed commentary on how different measurement variables influence the assessment of publication rates elsewhere [31].

We want to highlight that our study only assessed the extent of results reporting. We did not assess the reporting quality e.g. adherence to CONSORT [32]. We also did not assess whether identified papers reported all outcomes as specified in the registered protocol [1]. Finally, we did not compare whether results reported in published papers were consistent with results reported at the registry website [33, 34]. For judgments on the overall quality of clinical trial results reporting these other perspectives should be acknowledged as well.

In summary, the steady improvement in timely publication (within 24 months after CD) over time is promising. In contrast, the very low proportion of trials (7%) that report summary results in the registry is alarming, as most trials thus forego an important opportunity to increase their scientific and social value. Additionally, more recent trials might get published in a timely manner, but old trials still have relevant information that remains unavailable and unused. These results, which are both promising and alarming, should encourage German UMCs and other stakeholders, such as patient and funding organizations, to further improve their efforts and develop policies for the timely publication of trial results for future trials, as well as already finished yet unpublished trials. The publicly available Shiny app (http://s-quest.bihealth.org/intovalue/) might further be used to raise awareness about this element of good scientific practice in the scientific community and in the public.

## Supporting information

Supplement logistic regression

## Contributorship statement

DS, JM and UD designed the study. SW, NR, KW, HK, SO, CS, SK, UK and BS performed the search. NR, SK and BS performed the statistical analysis. All authors were involved in writing and editing the manuscript.

## Software availability

The R script used to as part of this study is available from https://doi.org/10.17605/OSF.IO/FH426 Archived source code at time of publication: https://doi.org/10.17605/OSF.IO/FH426

Licence: MIT License

## Data availability

All data underlying the results are available as part of the article and at OSF: Dataset 1: IntoValue. https://doi.org/10.17605/OSF.IO/FH426 ^15^

The data is available under a CC0 Licence. No additional source data are required.

## References

[1] Chan AW, Hrobjartsson A, Haahr MT, Gotzsche PC, Altman DG. Empirical evidence for selective reporting of outcomes in randomized trials: comparison of protocols to published articles. JAMA. 2004;291:2457–65.

[2] Hart B, Lundh A, Bero L. Effect of reporting bias on meta-analyses of drug trials: reanalysis of meta-analyses. BMJ. 2012;344:d7202.

[3] Kreis J, Panteli D, Busse R. How health technology assessment agencies address the issue of unpublished data. Int J Technol Assess Health Care. 2014;30:34–43.

[4] Mattina J, Carlisle B, Hachem Y, Fergusson D, Kimmelman J. Inefficiencies and Patient Burdens in the Development of the Targeted Cancer Drug Sorafenib: A Systematic Review. PLoS Biol. 2017;15:e2000487.

[5] Kimmelman J, London AJ. The structure of clinical translation: efficiency, information, and ethics. Hastings Cent Rep. 2015;45:27–39.

[6] Kiley R, Peatfield T, Hansen J, Reddington F. Data Sharing from Clinical Trials - A Research Funder’s Perspective. N Engl J Med. 2017;377:1990–2.

[7] Taichman DB, Backus J, Baethge C, Bauchner H, de Leeuw PW, Drazen JM, et al. Sharing Clinical Trial Data: A Proposal From the International Committee of Medical Journal Editors. JAMA. 2016;315:467–8.

[8] WHO. WHO Statement on Public Disclosure of Clinical Trial Results 2015. Available from: http://www.who.int/ictrp/results/reporting/en. 2015.

[9] Chalmers I. Underreporting research is scientific misconduct. JAMA. 1990;263:1405–8.

[10] Ross JS, Mulvey GK, Hines EM, Nissen SE, Krumholz HM. Trial publication after registration in ClinicalTrials.Gov: a cross-sectional analysis. PLoS Med. 2009;6:e1000144.

[11] Kasenda B, von Elm E, You J, Blumle A, Tomonaga Y, Saccilotto R, et al. Prevalence, characteristics, and publication of discontinued randomized trials. JAMA. 2014;311:1045–51.

[12] Krockenberger K, Bruns I, Ziegler A. [7th revision of the declaration of Helsinki: more than a recommendation?]. Dtsch Med Wochenschr. 2014;139:367–8.

[13] Chen R, Desai NR, Ross JS, Zhang W, Chau KH, Wayda B, et al. Publication and reporting of clinical trial results: cross sectional analysis across academic medical centers. BMJ. 2016;352:i637.

[14] Schmucker C, Schell LK, Portalupi S, Oeller P, Cabrera L, Bassler D, et al. Extent of non-publication in cohorts of studies approved by research ethics committees or included in trial registries. PLoS One. 2014;9:e114023.

[15] Dwan K, Gamble C, Williamson PR, Kirkham JJ, Reporting Bias G. Systematic review of the empirical evidence of study publication bias and outcome reporting bias - an updated review. PLoS One. 2013;8:e66844.

[16] De Angelis C, Drazen JM, Frizelle FA, Haug C, Hoey J, Horton R, et al. Clinical trial registration: a statement from the International Committee of Medical Journal Editors. N Engl J Med. 2004;351:1250–1.

[17] Zarin DA, Tse T. Medicine. Moving toward transparency of clinical trials. Science. 2008;319:1340–2.

[18] World Medical Association. Declaration of Helsinki: Ethical Principles for Medical Research Involving Human Subjects, Fortaleza. 2013.

[19] WHO. Joint statement on public disclosure of results from clinical trials; www.who.int/ictrp/results/jointstatement/en/. 2017.

[20] Begley CG, Buchan AM, Dirnagl U. Robust research: Institutions must do their part for reproducibility. Nature. 2015;525:25–7.

[21] Moore S, Neylon C, Paul Eve M, Paul O’Donnell D, Pattinson D. “Excellence R Us”: university research and the fetishisation of excellence. Palgrave Communications. 2017;3:16105.

[22] Goldacre B, DeVito NJ, Heneghan C, Irving F, Bacon S, Fleminger J, et al. Compliance with requirement to report results on the EU Clinical Trials Register: cohort study and web resource. BMJ. 2018;362:k3218.

[23] Powell-Smith A, Goldacre B. The TrialsTracker: Automated ongoing monitoring of failure to share clinical trial results by all major companies and research institutions. F1000Res. 2016;5:2629.

[24] ClinicalTrials.gov. Map of All Studies on ClinicalTrials.gov, https://clinicaltrials.gov/ct2/search/map?map=EU. accessed 2019.01.31.

[25] EUCTR. https://www.clinicaltrialsregister.eu/ctr-search/search. accessed 2019.01.31.

[26] ICMJE. http://www.icmje.org/about-icmje/faqs/clinical-trials-registration/. accessed 15 August 2018.

[27] Eaton L. UK universities show the way in reporting clinical trial results. BMJ. 2019;365:l1834.

[28] Wellcome Trust. Policy on clinical trials; 5. Post-trial requirements, https://wellcome.ac.uk/funding/guidance/wellcome-trust-policy-position-clinical-trials. accessed 2019.02.05.

[29] Blumle A, Meerpohl JJ, Schumacher M, von Elm E. Fate of clinical research studies after ethical approval--follow-up of study protocols until publication. PLoS One. 2014;9:e87184.

[30] Fleminger J, Goldacre B. Prevalence of clinical trial status discrepancies: A cross-sectional study of 10,492 trials registered on both ClinicalTrials.gov and the European Union Clinical Trials Register. PLoS One. 2018;13:e0193088.

[31] Strech D, Sievers S, Märschenz S, Riedel N, Wieschowski S, Meerpohl J, et al. Tracking the timely dissemination of clinical studies. Characteristics and impact of 10 tracking variables [version 1; referees: awaiting peer review], F1000Research. 2018;7.

[32] Turner L, Shamseer L, Altman DG, Schulz KF, Moher D. Does use of the CONSORT Statement impact the completeness of reporting of randomised controlled trials published in medical journals? A Cochrane review. Systematic reviews. 2012;1:60.

[33] Hartung DM, Zarin DA, Guise JM, McDonagh M, Paynter R, Helfand M. Reporting discrepancies between the ClinicalTrials.gov results database and peer-reviewed publications. Annals of internal medicine. 2014;160:477–83.

[34] Riveros C, Dechartres A, Perrodeau E, Haneef R, Boutron I, Ravaud P. Timing and completeness of trial results posted at ClinicalTrials.gov and published in journals. PLoS Med. 2013;10:e1001566; discussion e.

