## Supplement logistic regression for "Results dissemination from clinical trials conducted at German university medical centres was delayed and incomplete"

Method:

To identify variables with a strong effect on the probability of publication in an exploratory analysis, we used a logistic regression model to test several explanatory variables. The binary outcome of the logistic regression is timely publication (defined as publication < 24 month after the completion date, where publication can be a journal article or a summary result posted on the registry). We tested the following variables that were already defined in our preregistered protocol: industry (yes/no), medication (yes/no), primary completion years (2009-2013), phase (I-IV), institution size (small/large, threshold defined as median of the total number of registered trials per institution; institutions with a number of registered trials larger than the threshold are considered large, otherwise small), number of participants, and number of centers (mono-/multicentric).

To identify the relevant explanatory variables, the logistic regression model was built stepwise,

starting with univariate models and adding variables in each step. A log-likelihood ratio is used in each step to compare the more complex model (with an additional variable) to the less complex. In each step the variable that leads to the largest log-likelihood ratio is included in the subsequent model until adding subsequent variables did not further improve the model. As this is an exploratory analysis, we have not defined strict rules for variable inclusion a priori, not did we consider interaction terms here.

Results:

The center size can only reasonably be defined in the context of this study if one of the German UMCs is the leading center of the trial. Thus we first test a logistic regression model on the lead trials only, comparing a model with the center size as explanatory variable against the constant model. Both models lead to the same log likelihood rounded to two decimal places (-992.29). Therefore, we exclude the center size variable from the further analysis conducted on the full set of trials.

| Variable | Odds ratio (95% CI) | Log-likelihood score |
| --- | --- | --- |
| Constant | - | -1430.8 |
| Industry sponsor | 1.67 (1.40 – 2.01) | -1415.0 |
| Medication | 1.05 (0.89 – 1.26) | -1430.6 |
| Primary Completion Year (Odds ratio w.r.t. 2009) |  | -1426.4 |
| 2009 | 1 (ref) |  |
| 2010 | 1.08 (0.80 – 1.46) |  |
| 2011 | 1.05 (0.79 – 1.41) |  |
| 2012 | 1.17 (0.87 – 1.56) |  |
| 2013 | 1.51 (1.13 – 2.01) |  |
| Phase (Odds ratio w.r.t. Phase I) | 1 (ref) | -1423.7 |
| I-II | 1.43 (0.80 – 2.55) |  |
| II | 1.22 (0.78 – 1.91) |  |
| II-III | 0.93 (0.50 – 1.70) |  |
| III | 1.64 (1.05 – 2.57) |  |
| IV | 1.28 (0.80 – 2.05) |  |
| No phase (Non-drug trials) | 1.09 (0.71 – 1.67) |  |
| Number of participants | 1.00035 (1.0016 – 1.0055) (Increase in odds per participant, increase by 500 participants increases odds by 1.19) | -1422.8 |
| Multicentric study | 1.66 (1.39 – 1.98) | -1413.6 |

Table S1: Results for the univariate logistic regression models with each of the possible explanatory variables added to the model. The highest log-likelihood scores are reached by the variables “Mono/Multicentric” and “Industry sponsor”.

As can be seen in Table S1, for the univariate model the variables ‘Mono/Multicentric’ and ‘Industry sponsor’ where among the highest to increase the log-likelihood score. The third-highest log-likelihood is reached through the ‘number of participants’ variable. All three variables are strongly correlated. Multicenter studies tend to have higher participant numbers and tend to be industry sponsored (median enrollment: 137, 62% industry sponsored) compared to monocentric trials (median enrollment: 50, 11% industry sponsored).

Using the stepwise procedure, the following variables are added to the model: ‘Mono/Multicentric’ (resulting in log-lik. -1413.6, compared to -1430.8 for the constant model), ‘Industry sponsor’ (log-lik. -1409.1), ‘Primary completion year’ (log-lik. -1404.2), and ‘Number of participants’ (log-lik. -1399.6). While the strongest decrease in likelihood appears in the first step, the next three selection steps yield only a smaller decrease. Adding the next variable (i.e. the ‘phase’ variable) did not relevantly improve the log likelihood score (log-lik. -1397.8). The area under the curve, which is a measure for the predictiveness of the model, yields a value of AUC = 0.60 (95% CI 0.58 – 0.62) for the final model, which is a rough indication that the selected variables had only limited predictive power.

If we conduct the same analyses, but with timely publication defined as publication through a journal article alone, the only explanatory variable in the final model was the ‘number of participants’ variable. However, the predictive power is very limited in this case (AUC = 0.55).

Discussion:

The logistic regression together with a model selection procedure was used as an exploratory procedure to identify explanatory variables that have an effect on the publication probability of a clinical trial in our dataset.

The strongest effect on publication probability was seen for the variables ‘mono-/multicentric’, ‘industry sponsor’, and ‘number of participants’. Those variables are strongly correlated and thus can be seen as describing the same category of large, industry sponsored, multi-centric trials. This is not unexpected as these trials are costly in terms of both time and expenses and presumably more is dependent on the timely publication of these results. However, it needs to be kept in mind that the most important conclusion that can be drawn from our data, as can be seen by the low AUC-values, is that none of the tested variables is a very strong predictor for timely publication of results.

The data can be explored via our shiny app (<http://s-quest.bihealth.org/intovalue/>), but can also be downloaded for further analyses (see <https://osf.io/fh426/>)
